# Age variability and time averaging in oyster reef death assemblages

**DOI:** 10.1101/2022.11.05.515182

**Authors:** Stephen R. Durham, Gregory P. Dietl, Quan Hua, John C. Handley, Darrell Kaufman, Cheryl P. Clark

**Affiliations:** Florida Department of Environmental Protection, 2600 Blair Stone Road, MS 235, Tallahassee, FL, USA 32399; Paleontological Research Institution, Ithaca, NY, USA 14850; Department of Earth and Atmospheric Sciences, Cornell University, Ithaca, NY, USA 14853; Australian Nuclear Science and Technology Organisation, Locked Bag 2001, Kirrawee DC, NSW, Australia 2232; Simon School of Business, University of Rochester, Rochester, NY, USA 14627; School of Earth and Sustainability, Northern Arizona University, Flagstaff, AZ, USA 86011

## Abstract

A lack of temporal context for paleoecological data from molluscan death assemblages (DAs) makes integrating them with monitoring data from living communities to inform habitat management difficult. Here we illustrate this challenge by documenting the spatial and stratigraphic variability in age and time-averaging of oyster reef death assemblages. We radiocarbon dated a total of 573 oyster shells from samples of two burial depths on 28 oyster reefs around Florida and found 1) that spatial and stratigraphic variability in DA sample ages and time-averaging are of similar magnitude, and 2) that the shallow oyster reef DAs are among the youngest and highest-resolution molluscan DAs documented to-date, with most having time-averaging estimates of decades or less. This information increases the potential usefulness of the DAs for habitat management because measured indicators can be placed in temporal context relative to monitoring data. More broadly, the results highlight the potential to obtain decadal-scale resolution from oyster bioherms in the fossil record.

## INTRODUCTION

Decades of work on death assemblages (DAs) have successfully documented temporal changes in community composition or species attributes over time from direct assessments of the remains themselves (e.g., Kowalewski et al., 2000; Kidwell, 2007; Dietl and Durham, 2016; Albano et al., 2021), or proxy information derived from them (e.g., Gillikin et al., 2019). Despite the promise of these geohistorical records for conservation paleobiology, examples of their use by resource managers are still uncommon. One reason is the difficulty of putting DA data in temporal context. Geochronological analyses (e.g., radiocarbon dating) are expensive and difficult to interpret, leading many conservation paleobiological studies to either work around age-related uncertainties by citing general assumptions and/or studies from similar depositional settings (e.g., Dietl and Durham, 2016).

However, assemblage- or specimen-level chronological control is often required to meaningfully compare DA data with the annual or sub-annual real-time monitoring data typically used for resource management. This was the case for a project that was co-developed by the Florida Department of Environmental Protection (FDEP) Office of Resilience and Coastal Protection (ORCP) and the Paleontological Research Institution (PRI) where DA samples from oyster reefs were used to address a need for additional historical body size data on oyster populations for ORCP’s Statewide Ecosystem Assessment of Coastal and Aquatic Resources (SEACAR) project^1^.

Habitat management within the aquatic preserves managed by ORCP is conducted mainly with reference to conditions at the times they were established, which range from 1966 to 2020, meaning the ultimate utility of the DA approach to supplementing monitoring data for SEACAR depended on the specific age and time-averaging properties of the oyster reef DAs. We hypothesized that oyster reef structure might limit post-burial stratigraphic mixing enough that samples from the DAs could yield data at a high-enough temporal resolution to be integrated with real-time monitoring data from living oyster populations. To test this assumption and develop an understanding of both oyster reef taphonomy and the potential utility of DA data for FDEP, we produced a geochronological dataset to quantify the absolute ages and temporal resolutions of oyster reef DAs from around the state.

Here we describe this investigation and show that oyster reef DAs preserve reliably recent and high-resolution stratigraphic records relative to most other molluscan DAs documented to-date. The oyster reef DAs also had some of the lowest estimated scales of time-averaging of any molluscan DA, suggesting these records are often appropriate for decadal-scale conservation paleobiological investigations. We also highlight the geographic variability in our dataset and its implications for the importance of location-specific geochronological information for increasing the salience of paleoecological data for the resource management community.

## MATERIAL AND METHODS

In order to build a geochronological dataset to evaluate the utility of oyster DA samples for documenting trends over recent decades, a total of 573 *C. virginica* left valve specimens was randomly selected from oyster DA samples representing two stratigraphic intervals (15-25 cm and 25-35 cm) from up to three sample holes at each of 28 oyster reefs in 10 locations around Florida, i.e., between 2 and 7 specimens from each DA sample (Fig. 1). The selected specimens^2^ were dated by radiocarbon analysis of powdered carbonate targets (Bush et al., 2013; Hua et al., 2019)—a less expensive method with lower precision than the standard analysis of graphite targets, but which yields similar ages (Bright et al., 2021)—to achieve a higher sample size. Specimens were prepared at Northern Arizona University and analyzed at the W.M. Keck Carbon Cycle Accelerator Mass Spectrometry facility at the University of California, Irvine. Local corrections for the hardwater effect (e.g., Spennemann and Head, 1998) and/or estuarine influences (e.g., Ulm et al., 2009), in terms of dead carbon contribution, were developed using additional radiocarbon analyses of 1-2 live-caught oyster specimens from each sampling area (Supplementary Information).

**Figure 1.**
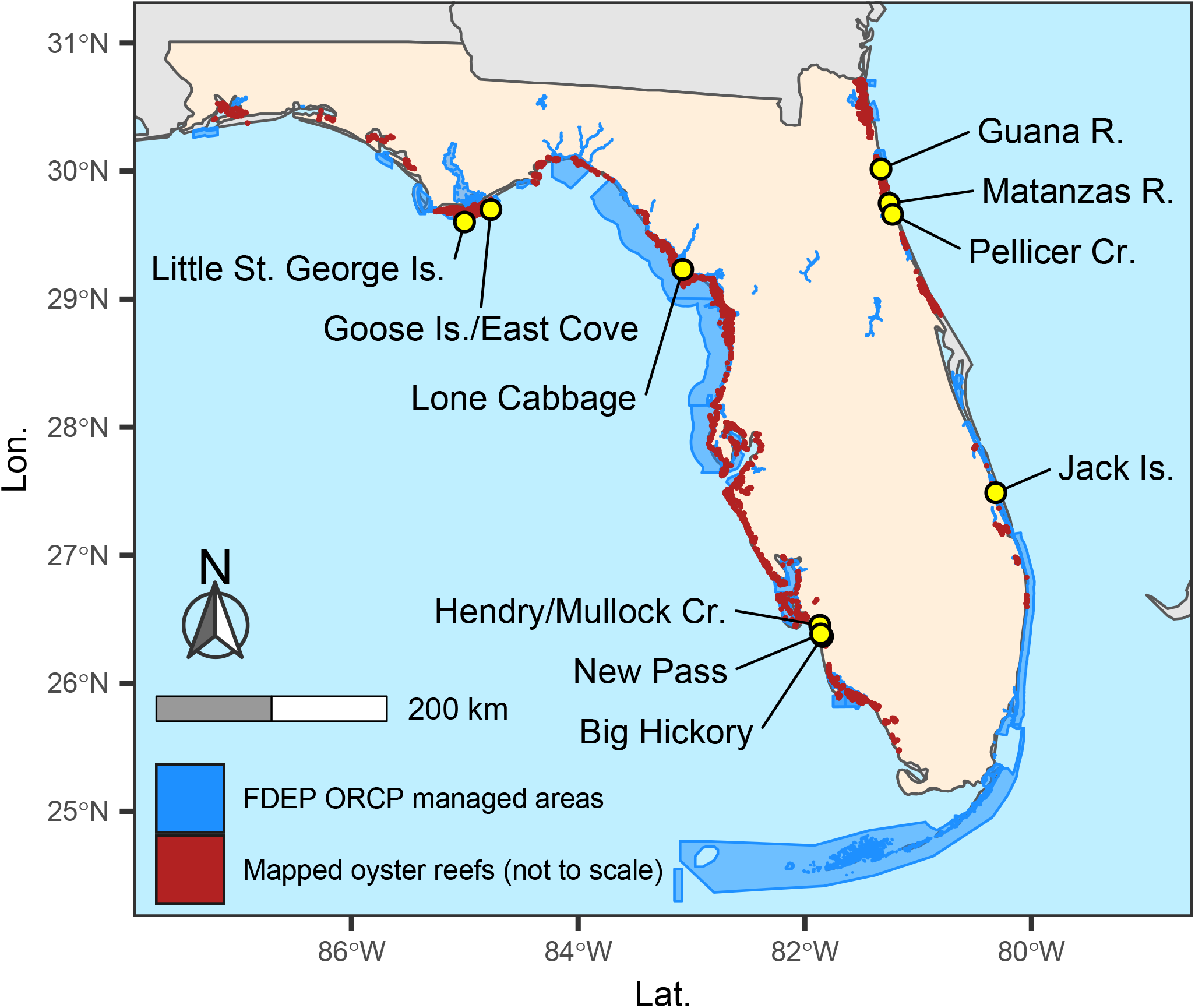
Map showing the 10 localities in Florida where oyster reef DAs were sampled (yellow circles). FDEP ORCP = Florida Department of Environmental Protection Office of Resilience and Coastal Protection.

Age calibration was performed using OxCal v4.4 software (Bronk Ramsey, 2009), and the Marine20 calibration curve (Heaton et al., 2020) with a constant regional marine reservoir correction—ΔR = −134 ± 26 years, which is equivalent to 5 ± 32 years (Kowalewski et al., 2018) relative to Marine13 (Reimer et al., 2013)—extended to 2019 using an updated version of the regional marine bomb radiocarbon data presented in Kowalewski et al. (2018) (Supplementary Information). Following Kowalewski et al. (2018), we used empirical posterior distributions of age probabilities for the specimens in each DA sample to estimate 1) DA sample age^3^ as the median of the specimen ages weighted by their probabilities, 2) the total age variability in the DA sample as the interquartile range of the specimen ages weighted by their probabilities (IQR_TAV_) and 3) the age-estimation error for individual specimens in the DA sample as the median of the interquartile ranges of the specimen ages, weighted by their probabilities. The difference between the IQR_TAV_ and the specimen age estimation error for a given DA sample—i.e., the corrected posterior age estimate (CPE, *sensu* Kowalewski et al., 2018; also known as residual time averaging in some studies)—is an estimate of the time averaging in the sample, accounting for specimen age estimation error.

To compare the contributions of location and burial depth to overall variation in DA sample median age and CPE, we fit a hierarchical Bayesian model to the data for each burial depth as well as the burial depth difference for each DA sample hole (Supplementary Information). All data analyses were conducted using R statistical software v4.1.1 (R Core Team, 2021) in the RStudio integrated development environment (RStudio Team, 2021).

## RESULTS

The radiocarbon results indicated that oyster reef DAs are high-resolution archives with abundant shells from the recent past and minimal time-averaging. Among the 114 dated oyster DA samples, median calibrated ages ranged from 1622 to 2014, but 93.9 % were post-1950 (Fig. 2), and 53.5 % of the DA samples had sub-decadal-scale CPE (0-10 years), 36.8 % had decadal-scale CPE (11-100 years), and 9.6 % had centennial-scale CPE (101-1000 years) (Fig. 3; see Appendix DR1 for DA sample-level results). Moreover, co-located samples from different burial depths showed the expected temporal order (i.e., deeper = older) in most cases: out of the 49 sample holes for which both depth intervals were processed and dated, 10 had median DA sample ages for the 15-25 cm burial depth that were older than those of the 25-35 cm burial depth material, and six of those cases were from a single locality (Fig. 2). The results also showed that the age and time-averaging of a given burial depth can vary substantially over small spatial scales (i.e., both intra- and inter-reef assemblage variation; Fig. 2). In fact, the modeled standard deviations (SD) for spatial variability in median age and CPE (e.g., 53.1 and 53.9 years for the 15-25cm depth, DA sample-hole-level median SDs for median age and CPE, respectively) were of similar magnitude to those for the difference between burial depths (e.g., 58.1 and 96.2 years for the DA sample-hole-level median depth differences SDs for median age and CPE, respectively; Table DR4, Figures DR2 to DR7).

**Figure 2.**
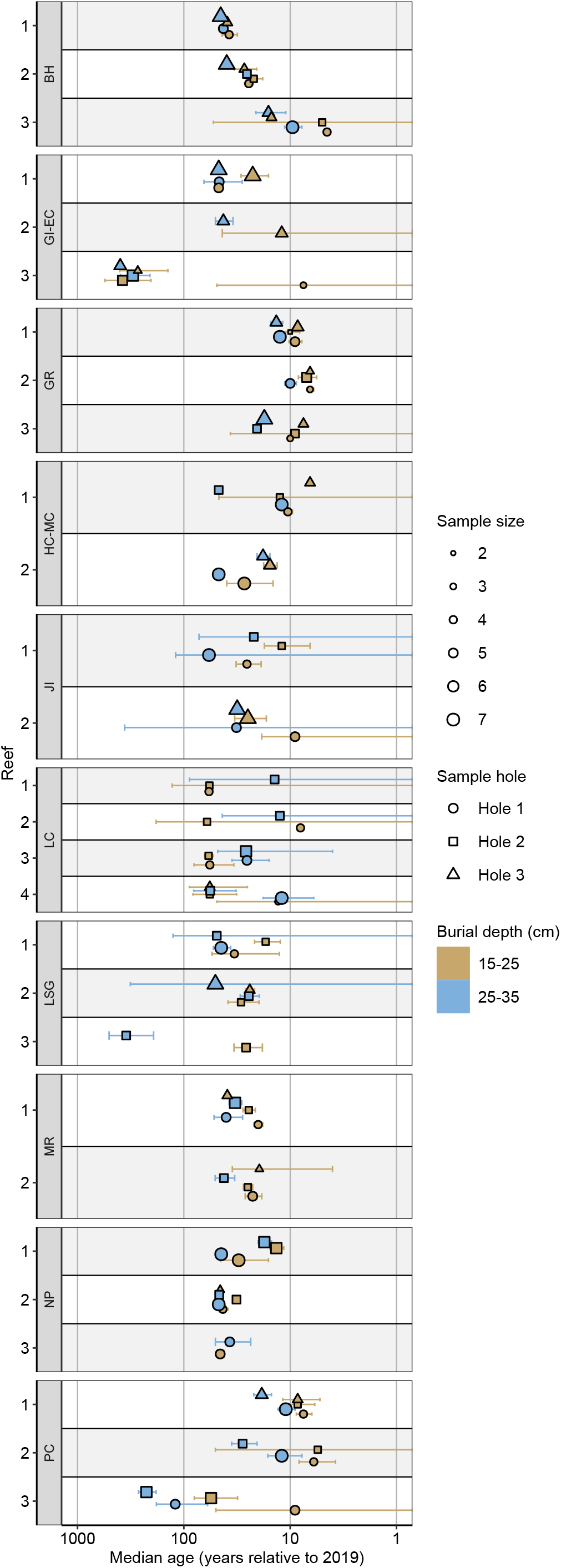
Plot showing the median ages of the oyster DA samples by reef and locality relative to 2019. Note that the x-axis is on the log10 scale. Error bars represent the corrected posterior age estimate for each bulk sample. Localities are listed on the y axis in counter-clockwise geographic order around the state, starting at the panhandle: LSG = Little St. George Island, GI-EC = Goose Island/East Cove, LC = Lone Cabbage, HC-MC = Hendry Creek/Mullock Creek, NP = New Pass, BH = Big Hickory, JI = Jack Island, PC = Pellicer Creek, MR = Matanzas River, GR = Guana River.

**Figure 3.**
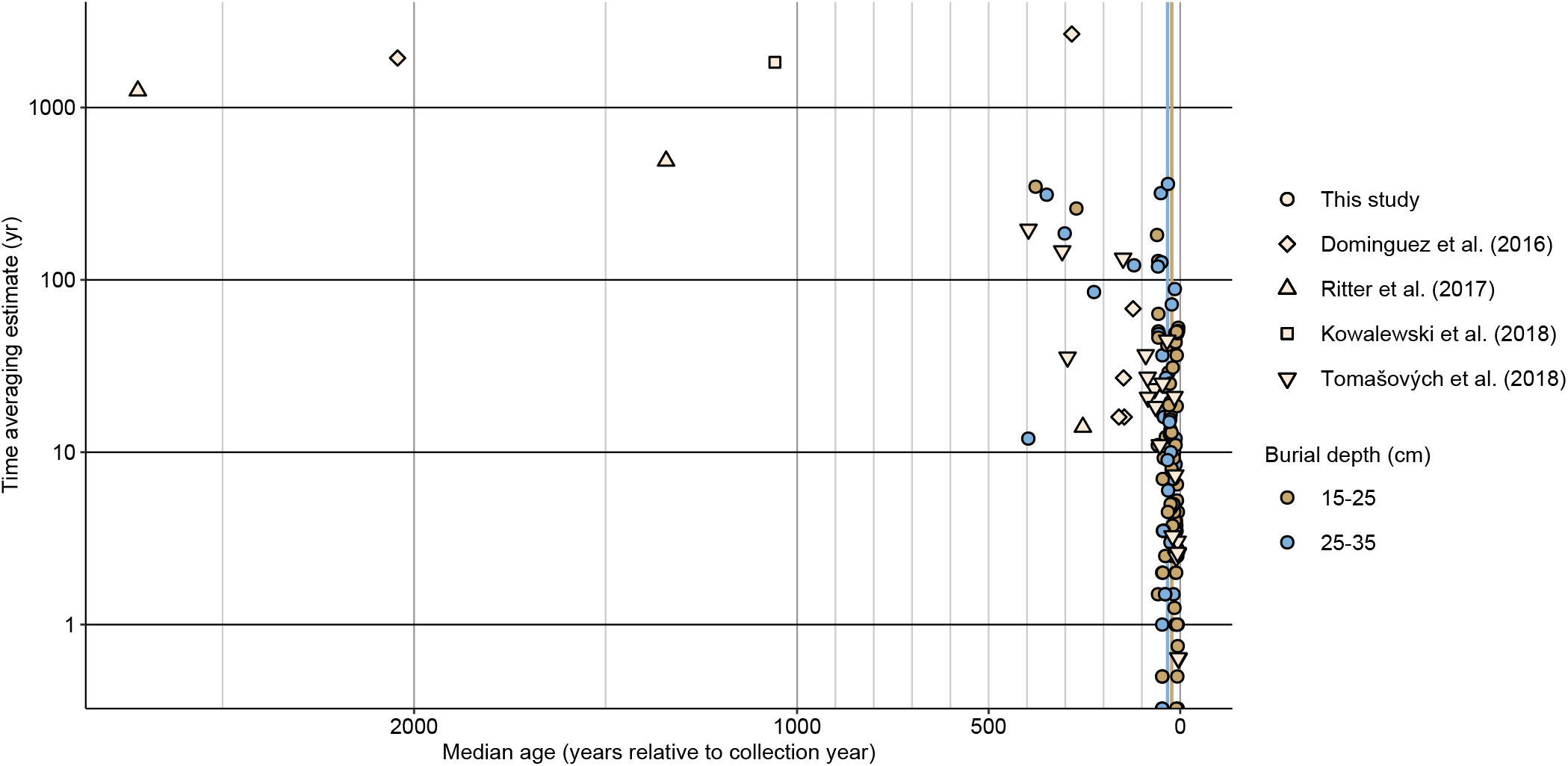
Plot of median ages (relative to collection year) against corrected posterior age estimates for some comparable recent studies of molluscan DAs. The *C. virginica* DA samples are consistently younger and, in many cases, less time-averaged than other studied DAs. Thin brown and blue vertical lines show the medians of the median ages from this study for the DA samples from 15-25 cm and 25-35 cm burial depths, respectively. Note the y-axis is shown on the log10 scale.

## DISCUSSION

To our knowledge, this is the largest study of age-depth relationships, and the first to document time-averaging, in oyster reef DAs. We found that relative to other molluscan DAs, the oyster DA samples were younger, less time-averaged, and had less spatial variability in both calibrated age and time-averaging estimates (Flessa et al., 1993; Meldahl et al., 1997; Kowalewski et al., 1998, 2018; Kosnik et al., 2009, 2015; Krause et al., 2010; Dexter et al., 2014; Dominguez et al., 2016; Ritter et al., 2017; Tomašových et al., 2019; Albano et al., 2020; see additional studies summarized in Table 1 of Kidwell, 2013; but see also Tomašových et al., 2018 for an example of a non-reef DA with decadal-scale resolution).

Among the few recent studies that have used similar methods to quantify scales of time-averaging, the *C. virginica* DA samples typically had younger median ages by about 100 years and over half of our samples also had lower CPE, some by an order of magnitude or more (Fig. 3). For instance, Dominguez et al. (2016) sampled the upper 20 cm of sediment at six sites with ~9 m water depth in Sydney Harbour, Australia and found decadal scales of time averaging (~20-40 years) in DAs of the bivalve *Fulvia tenuicostata*, but the median ages of the samples were ~150 years. In contrast, median CPE across all of the *C. virginica* DA samples in our study was ~9 years (ranging from zero to 360 years) and the medians of the median calibrated ages across all of the 15-25 cm and 25-35 cm burial depth DA samples were 22 years and 33 years, respectively. The SDs for both median age and CPE among the locations sampled by Dominguez et al. (2016) were both higher than the respective modeled locality-level SDs for the oyster reefs we sampled, despite the much greater geographic area covered by our study (Table DR3, Figures DR2 to DR7).

One exception to this pattern is Tomašových et al. (2018), who found comparable age and time-averaging estimates to ours in *Corbula gibba* DAs from cores of the Po and Isonzo prodeltas (Fig. 3). However, the authors stated that the two deltas have some of the highest sedimentation rates in the northern Adriatic Sea, and median ages and time-averaging estimates for *C. gibba* DAs from the eastern Gulf of Trieste—across the Gulf from the Isonzo River and characterized by low sedimentation rates—were older and more time-averaged than the prodelta samples by nearly two orders of magnitude (Tomašových et al., 2019). This large difference between the two depositional settings suggests that high resolution is not a general characteristic of *C. gibba* DAs in the region. In contrast, decadal-scale resolution appears to be a common feature of DAs from intertidal *C. virginica* reefs in multiple estuaries across Florida.

Overall, our results suggest oyster reefs have a relatively high shell burial rate and less stratigraphic mixing relative to non-reef molluscan DAs, supporting the hypothesis that the physical structure of oyster reefs limits their DAs’ susceptibility to some taphonomic processes. Despite their higher temporal resolution than other types of molluscan DAs, however, considerable geographic variation, and even intra-assemblage variation, was still present in the oyster DA median ages and CPE (Fig. 2), precluding useful generalizations of the results into regional or statewide guidance on age vs. burial depth relationships or scales of time-averaging (see Supplementary Information for an example).

This variability illustrates why specific geochronological information will be important for many conservation paleobiological contributions to oyster management. Exactly how necessary they are for any given project will depend on the questions being investigated, but trends in many indicators of oyster population condition, such as live oyster size-frequency, are typically tracked at annual or sub-annual intervals by oyster monitoring programs. To integrate measurements from DA samples with such high-resolution records for trend analyses, it will likely be necessary to know, for instance, whether the median calibrated age and CPE of a DA sample are 2002 and 9.25 years, respectively, or 1986 and 41.75 years—as was the case for two of the 15-25cm burial depth DA samples from our Little St. George Island locality.

Once these data are obtained and DA sample ages and time-averaging can be estimated, however, more confident comparisons between the DA data and monitoring data become possible, such as integrating DA and real-time data into a single model that accounts for uncertainty in sample ages, instead of only focusing on more general “before/after” comparisons (e.g., Dietl and Durham, 2016). Specifically, the limited time-averaging and recent median ages make these oyster DA samples a promising resource for decadal-scale historical oyster information for the ORCP SEACAR project. Most of the DA samples represent a relevant time period for management and could yield historical oyster population data for ORCP managed areas that are not otherwise accessible, given the lack of long-term oyster monitoring records from most coastal areas of the state.

Lastly, given the apparently limited stratigraphic mobility of shells preserved within Recent oyster DAs and the fact that oyster reefs are sometimes preserved in the fossil record as *in situ* bioherms, our study results suggest the intriguing possibility that the degree of time-averaging in a fossil bioherm is not dramatically greater than in the DA of a living oyster reef. If this is the case, bioherms may preserve decadal-scale records from time-periods when information at such a fine temporal resolution is exceptionally rare, making them potentially valuable records for studies of short-term ecological processes in the deep past that are not possible with other fossil assemblages (e.g., Kowalewski et al., 1998; Kidwell and Tomasovych, 2013).

## ACKNOWLEDGMENTS

We thank Jordon Bright (Amino Acid Geochronology Lab, Northern Arizona University) and the Keck AMS lab (University of California, Irvine) for ^14^C analyses, and Matthew Kosnik (Macquarie University) for help adapting the Kowalewski et al. (2018) R script. We also thank colleagues and volunteers from FDEP, PRI, and Florida Department of Agriculture and Consumer Services (FDACS) who helped with field or lab work (*FDEP, ^†^PRI, ^‡^FDACS, ‘Pelican Island National Wildlife Refuge). <u>Staff</u>: M. Anderson*, P. Benjasirichai^‡^, E. Bourque*, M. Brown*, C. Brunk*, C. Clark^‡^, S. Cofone^‡^, R. Cray*, E. Dark*, M. DeHaven^‡^, N. Dix*, S. Erickson*, J. Fleiger^‡^, J. Garwood*, T. Green*, B. Hamill*, K. Harshaw^‡^, D. Hersl*, T. Jones*, K. Lang*, P. Marcum*, M. McMurray*, B. Mowbray*, R. Noyes*, J. Pier^†^, R. Prado*, M. Pruden^†^; <u>Volunteers</u>: B. Alexander*, A. Bishop*, S. Gavirneni^†^, C. Hormuth*, A. McNeil^†^, M. Melekos^†^, R. Mondazzi*, D. Philipp*, E. Prest^†^, M. Raymond*, M. Schilling*, Z. Siper^†^, B. Skoblick^†^, A. Thorsness*, D. Thorsness*, J. Valentine^•^. We also gratefully acknowledge funding from the National Oceanic and Atmospheric Administration (NOAA) Office for Coastal Management under the Coastal Zone Management Act of 1972, as amended, to the Florida Coastal Management Program (NOAA awards NA18NOS4190080 and NA19NOS4190064). The views, statements, findings, conclusions and recommendations expressed herein are those of the author(s) and do not necessarily reflect the views of the State of Florida, NOAA or any of their subagencies.

## Footnotes

1 www.floridadep.gov/SEACAR

2 See Durham et al. (2019) for DA sample collection and processing information and Supplementary Information for details on specimen selection for radiocarbon analysis.

3 We use the terms “specimen age” and “sample age” to refer to radiocarbon results for an individual oyster shell and all oyster shells from a given DA sample, respectively.

## Notes

https://data.florida-seacar.org/programs/details/5035

### Competing Interest Statement

The authors have declared no competing interest.

