## Supplementary Information for "Age variability and time averaging in oyster reef death assemblages"

### DETAILS ON SAMPLING AND AMS RADIOCARBON ANALYSES

We focused our oyster reef sampling on 10 different areas around Florida that corresponded with Florida Department of Environmental Protection (FDEP) Office of Resilience and Coastal Protection (ORCP) managed areas and existing live oyster population monitoring data (Table S1). Using ArcGIS, 12 candidate reefs were randomly selected in each sampling area from the "Oyster_Beds_in_Florida" map layer[1], which was produced by the Oyster Integrated

[1] current version available from https://geodata.myfwc.com/datasets/oyster-beds-in-florida?geometry=-100.316%2C24.682%2C-66.939%2C31.461, accessed 5/19/2021.

Mapping and Monitoring Program of the Florida Fish and Wildlife Conservation Commission (Radabaugh, 2019). The map layer is not comprehensive, but it is the most complete Florida oyster reef map available. In the field, up to five reefs were semi-randomly selected for sampling from the 12 candidate reefs (or in some cases, nearby reefs) based on factors such as accessibility, tide level, and reef condition (e.g., presence of live oysters, likelihood of having a substantial death assemblage).

TABLE S1. SAMPLING LOCALITIES AND ORCP MANAGED AREAS

| ORCP Managed Area | Water Body | Region | Date of Designation | Area (acres) | Study Localities |
|---|---|---|---|---|---|
| Apalachicola Bay Aquatic Preserve | Apalachicola Bay | NW | 1969 | 80,876 | Little St. George Is. |
| Apalachicola National Estuarine Research Reserve | Apalachicola River, Apalachicola Bay | NW | 1979 | 234,691 | Little St. George Is., Goose Island/East Cove |
| Big Bend Seagrasses Aquatic Preserve | Gulf of Mexico (Apalachee Bay to Waccasassa Bay) | NW | 1985 | 984,325 | Lone Cabbage |
| Estero Bay Aquatic Preserve | Estero Bay | SW | 1966/1983 | 13,829 | Hendry Creek/Mullock Creek,<br>New Pass,<br>Big Hickory |
| Indian River-Vero Beach to Ft. Pierce Aquatic Preserve | Indian River Lagoon | NE | 1970 | 9,477 | Jack Island |
| Guana Tolomato Matanzas National Estuarine Research Reserve | Guana River, Tolomato River, Matanzas River, Pellicer Creek | NE | 1999 | 76,760 | Guana River,<br>Matanzas River,<br>Pellicer Creek |
| Guana River Marsh Aquatic Preserve | Guana River, Tolomato River, Atlantic Ocean (Ponte Vedra Beach, from Sawgrass to ~1.5km north of Vilano Beach) | NE | 1985 | 37,048 | Guana River |

Death assemblage (DA) sampling of each selected oyster reef was integrated with intertidal oyster reef monitoring methods used by FDEP staff (Dix and Marcum, 2018): a 30 m transect tape was extended parallel to the long axis of the reef and across the portion of the reef that appeared to have the densest accumulation of oysters, and three 0.0625 $m^2$ (25 cm x 25 cm) quadrats were placed at distances along the transect selected with a random number generator. At each quadrat, the top 15 cm of material was removed and placed to the side in order to reach a depth below the living oysters and at which buried shells were unlikely to be re-exhumed (i.e., below the taphonomically active zone; Powell et al., 2012; Rodriguez et al., 2014; Dix and Marcum, 2018). Once the hole was prepared, two DA samples were extracted comprising the subsequent two 10 cm depth intervals[2] (i.e., 15-25 cm and 25-35 cm below the reef surface). Each sample was collected into a 4 mil polyethylene sample bag labeled with the sample information. The 10 sampling areas were visited by the research team over the course of three field trips in late summer to fall of 2018; samples for each trip were cushioned with packing paper and sealed in moving boxes in groups of two to four before being transported to a climate-controlled (non-refrigerated) storage facility. After all fieldwork was completed, the boxed samples were transported to the Paleontological Research Institution in Ithaca, New York for processing and curation. Sampling was authorized by Environmental Resource Program Permit Exemption Verification 0366243-001-EE/19 (Florida Dept. of Environmental Protection), Special Activity License SAL-18-2064-SR (Florida Fish and Wildlife Conservation Commission), Division of Recreation and Parks Scientific Research/Collecting Permit 07051810 (Florida Department of Environmental Protection), Nationwide Permit Number 4 SAJ-2018-01876 (U. S. Army Corps of Engineers), and a Visiting Investigator Permit from the Guana Tolomato Matanzas National Estuarine Research Reserve, all issued to S. Durham. In addition, the target sampling areas were modified prior to beginning fieldwork in response to a review by

[2] The only exception was reef 1 from New Pass, for which 15-30cm and 30-45cm depth intervals were collected. The results for these samples were similar to those of the other New Pass reefs, so we did not distinguish between the 15-30cm and 15-25cm or the 30-45cm and 25-35cm DA sample results in our analysis.

the Florida Department of State Division of Historical Resources to ensure no impact to archaeological resources (DHR Project File 2018-3543). Ownership of all samples collected was transferred to the Paleontological Research Institution for long-term storage (PRI Accession Number 1860).

In the laboratory, each sample bag was emptied over stacked 6 mm and 1.9 mm mesh sieves, and a subsample of the matrix material was collected before the sediment was washed and the oyster shells were separated from the other material. All left valves ≥ 25 mm in shell height and estimated to be at least 90% complete in each DA sample were assigned numbers that were used to randomly select specimens for radiocarbon analysis and index specimens for additional data collection. Initially, 25 specimens were randomly selected for radiocarbon analysis from all numbered specimens across all processed DA samples from the same reef x stratigraphic interval (i.e., 15-25 cm or 25-35 cm burial depths). From those 25 specimens, 12-14 specimens were selected for analysis such that each processed DA sample was represented by at least two specimens. Otherwise, specimens were evaluated in the order in which they were selected and any specimens with substantial bioerosion or other damage to the hinge plate were rejected due to the higher potential for chemical alteration of the shell interior. However, it became clear early on that specimens from the same burial depth but different locations on a reef often varied in age, so we began randomly selecting 8-10 specimens from each processed DA sample, from which between four and seven specimens were chosen for analysis as previously described. This method ensured that most processed samples were represented by at least four specimens and that all reef x stratigraphic intervals were represented by at least five specimens.

A wedge of shell was cut out of the hinge plate of each selected specimen using a Gryphon C-40 diamond bandsaw, after which the fragments were air-dried at room temperature and placed in labeled polyethylene bags. All specimens were shipped to Northern Arizona University, where subsamples of the foliated calcite portions of each fragment were prepared for radiocarbon analysis following procedures modified from Bush et al. (2013). Briefly, fragments were leached in 2N HCl to remove approximately 30% of their mass, dried, and ground to a fine powder. Between 0.3 and 0.5 mg of carbonate was mixed with metal powder and pressed into accelerator mass spectrometry (AMS) targets. Once prepared, samples were analyzed at the W. M. Keck Carbon Cycle AMS facility at the University of California, Irvine. In order to date as many specimens as possible, the majority of analyses were performed on powdered carbonate targets, which are less costly to analyze, but have lower precision than the graphite targets used in standard AMS radiocarbon analyses (Bush et al., 2013; Hua et al., 2019; Bright et al., 2021). Eleven specimens were re-analyzed by standard AMS radiocarbon analyses to check the lower-precision radiocarbon results. Additional standard AMS analyses were also conducted on live-caught filter-feeding clams from near the mouth of Alligator Harbor, Florida (approximate lat./long.: 29.910016, -84.429683) and at least one live-caught oyster from each locality. These analyses were used to estimate local "dead carbon" corrections for the dead shell radiocarbon results before they were calibrated to calendar ages. The dead carbon contribution is likely due to the hardwater effect as a result of geological settings (e.g., Spennemann and Head, 1998) and/or estuarine influences resulting from riverine discharges with incomplete $^{14}C$ mixing with the open ocean (e.g., Ulm et al., 2009), and is assumed to affect all specimens within a site/reef equally through time. In addition, we dated four museum oyster specimens of known age to test the accuracy of their calibrated dates using our method of dead carbon corrections and age calibration. Live oyster specimens were collected either by staff from the Florida Department of Agriculture and Consumer Services under the Department's public health authority for the sanitary control of shellfish (*Florida Statutes* section 597.020 and Florida Administrative Code Rule 5L-1) or by FDEP staff under Special Activity License SAL-20-2259A-SR issued by the Florida Fish and Wildlife Conservation Commission to S. Durham. Ownership of all live-caught specimens was transferred to the Paleontological Research Institution for long-term storage (PRI Accession Number 1897).

### RADIOCARBON GEOCHRONOLOGY CALIBRATION AND RESULTS

Reduced major axis regression of eleven DA specimens by both low-precision and standard AMS showed a strong relationship (slope = 0.994, $R^2$ = 0.97), justifying our use of the lower-precision method (Fig. DR1). These results are similar to those of a much larger comparison study demonstrating the strong correspondence between radiocarbon values measured by the two AMS methods (Bright et al., 2021).

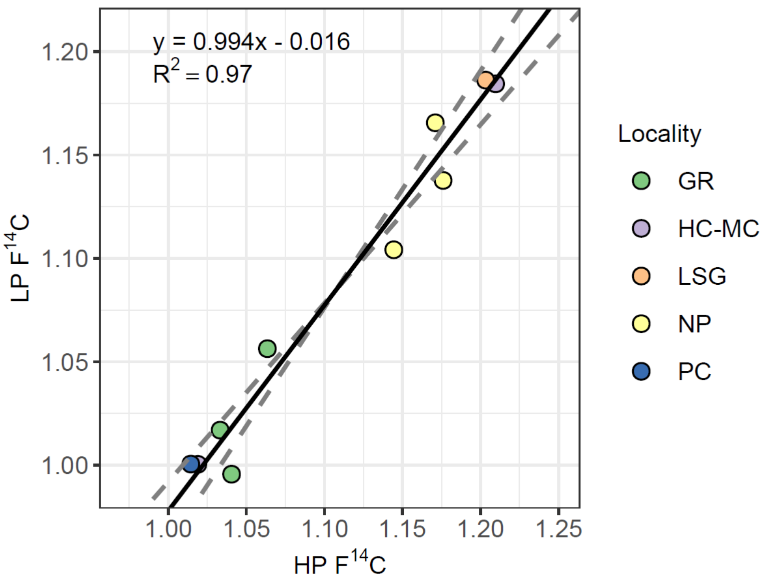


Figure DR1. Reduced axis regression of standard (high precision, graphite targets) AMS radiocarbon results in fraction modern carbon (HP $F^{14}C$) for death assemblage specimens against low-precision (carbonate target) AMS radiocarbon results (LP $F^{14}C$) for the same specimens. Solid line = reduced axis regression line, dashed lines = 97.5 % confidence interval. Localities are listed in counter-clockwise geographic order around the state, starting at the panhandle: LSG = Little St. George Island, HC-MC = Hendry Creek/Mullock Creek, NP = New Pass, PC = Pellicer Creek, GR = Guana River.

“Dead carbon” contribution in each sample area ($DeadC_{Local}$) was estimated using the fraction modern carbon (n = 1) or weighted mean fraction modern carbon (n = 2) value of local live-caught oyster specimens ($F^{14}C_{Oyster}$) and the weighted mean value of two live-caught *Mercenaria* sp. specimens ($F^{14}C_{Clam}$) from near the mouth of Alligator Harbor in northwest Florida (Tables DR1, DR2):

$$DeadC_{Local} = 1 - \frac{F^{14}C_{Oyster}}{F^{14}C_{Clam}} \qquad (1)$$

The clams were used because they came from a full-marine salinity environment, so were not expected to be influenced as much by hardwater and/or estuarine effects as the estuarine *C. virginica* specimens.

Prior to calibration, the $F^{14}C$ value for each dead shell specimen dated from each locality was corrected using the corresponding local dead carbon estimate:

$$F^{14}C_{Corrected} = \frac{F^{14}C_{Local}}{(1-DeadC_{Local})} \qquad (2)$$

where $F^{14}C_{Local}$ is the measured $^{14}C$ content of a dead shell specimen. The dead carbon corrected $F^{14}C$ standard deviations were calculated as:

$$SD_{Corrected} = \sqrt{\left(\frac{SD_{LocalF^{14}C}}{(1-DeadC_{Local})}\right)^2 + \left(SD_{LocalDeadC} \times \frac{F^{14}C_{Local}}{(1-DeadC_{Local})^2}\right)^2} \qquad (3)$$

where $SD_{LocalDeadC}$ and $SD_{LocalF14C}$ are uncertainties associated with the local dead carbon contribution and the measured $F^{14}C$ value of a dead shell specimen, respectively.

The corrected $F^{14}C$ values were then calibrated using OxCal v4.4 (Bronk Ramsey, 2009), and the Marine20 calibration curve (Heaton et al., 2020) with a constant regional marine reservoir correction—ΔR = -134 ± 26 years, which is equivalent to 5 ± 32 years (Kowalewski et al., 2018) relative to Marine13 (Reimer et al., 2013)—extended to 2019 using the regional marine bomb radiocarbon data (Kowalewski et al., 2018) and the weighted mean $F^{14}C$ value from the two live-caught *Mercenaria* sp. specimens from this study. The calibrated ages and time-averaging estimates for the oyster DA samples were calculated as described in the main text and Kowalewski et al. (2018). See Appendix DR1 for the DA sample ages and time-averaging estimates, Appendix DR2 for uncalibrated high-precision AMS radiocarbon results, Appendix DR3 for uncalibrated low-precision AMS radiocarbon results, and Appendix DR4 for the posterior probability distributions from the OxCal output for all calibrated radiocarbon ages.

TABLE DR2. DEAD CARBON ESTIMATES FOR EACH LOCALITY

| Locality | Genus | N | Weighted Mean $F^{14}C$ | ± | Dead C ($F^{14}C$) | ± |
|---|---|---|---|---|---|---|
| Little St. George Is. | *Crassostrea* | 2 | 1.0026 | 0.0069 | 0.0079 | 0.0071 |
| Goose Island/East Cove | *Crassostrea* | 2 | 1.0120 | 0.0014 | -0.0014 | 0.0022 |
| Alligator Harbor* | *Mercenaria* | 2 | 1.0106 | 0.0017 | NA | NA |
| Lone Cabbage | *Crassostrea* | 2 | 0.9225 | 0.0039 | 0.0872 | 0.0041 |
| Hendry Creek/Mullock Creek | *Crassostrea* | 2 | 0.9768 | 0.0030 | 0.0334 | 0.0034 |
| New Pass | *Crassostrea* | 2 | 0.9972 | 0.0042 | 0.0132 | 0.0045 |
| Big Hickory | *Crassostrea* | 1 | 0.9900 | 0.0022 | 0.0204 | 0.0027 |
| Jack Island | *Crassostrea* | 2 | 1.0082 | 0.0021 | 0.0024 | 0.0027 |
| Pellicer Creek | *Crassostrea* | 2 | 1.0109 | 0.0039 | -0.0003 | 0.0042 |
| Matanzas River | *Crassostrea* | 2 | 1.0229 | 0.0014 | -0.0122 | 0.0022 |
| Guana River | *Crassostrea* | 2 | 0.9974 | 0.0038 | 0.0130 | 0.0041 |

*Location for full-marine salinity clam specimens; no oysters collected.

[†]Localities are listed in counter-clockwise geographic order around the state, starting at the panhandle.

We also dated four specimens of known age from the Florida Museum of Natural History as a check on the dead carbon correction and age calibration procedures. Two of the four specimens had a known collection date of 1979 and were collected from the northwest coast of Cedar Key Island, within about 10 km of the Lone Cabbage locality. Using the local dead carbon correction developed for the Lone Cabbage locality, the median calibrated ages of the two museum specimens were both 1972 and their age ranges at 95% CI were 1967.0-1982.5 (Table DR3). The two other museum specimens were collected farther from our localities; one was collected in Indian Pass, Franklin County in 1938 (approximately 20 km northwest of the Little St. George Island locality) and the other was collected in Gordon Pass, Collier County in 1932 (approximately 30 km south of the Big Hickory locality), so the dead carbon corrections for those localities are not as likely to be appropriate as the Lone Cabbage values were for the Cedar Key Island specimens. Further, these specimens lived prior to the atmospheric nuclear testing in the 1950s and 1960s, which produced the "bomb pulse" radiocarbon signature that allows for much higher-resolution radiocarbon dating for many materials generated after the mid-1950s (Hua, 2009). Considering these factors, the median calibrated age of 1916.5 for the specimen from Gordon Pass was reasonable, while the broad calibrated age range associated with the median calibrated age of 1853.5 for the specimen from Indian Pass was not surprising. Altogether, the results of these analyses supported the validity of the age estimates from the dead carbon correction and radiocarbon calibration procedures.

TABLE DR3. RADIOCARBON RESULTS FROM SPECIMENS OF KNOWN AGE AND MINIMUM, MAXIMUM, AND MEDIAN CALIBRATED POSTERIOR AGES FOR EACH SPECIMEN

| FLMNH Cat. No.* | Collection Date | Nearest Locality[†] | Sample ID | $F^{14}C$ | ± | Dead C | ± | Corrected $F^{14}C$ | ± | Age Range Min. | Age Range Max. | Median Cal. Age |
|---|---|---|---|---|---|---|---|---|---|---|---|---|
| UF 15484 | 1938 | Little St. George Is. | UAL19523 | 0.9338 | 0.0018 | 0.0079 | 0.0071 | 0.9412 | 0.0069 | 1515.0 | 1961.0 | 1853.5 |
| UF 512435 | 1979 | Lone Cabbage | UAL19520 | 1.1573 | 0.0022 | 0.0872 | 0.0041 | 1.2678 | 0.0062 | 1967.0 | 1982.5 | 1972.0 |
| | | | UAL19521 | 1.1068 | 0.0024 | 0.0872 | 0.0041 | 1.2125 | 0.0061 | 1967.0 | 1982.5 | 1972.0 |
| UF 15491 | 1932 | Big Hickory | UAL19522 | 0.9405 | 0.0019 | 0.0250 | 0.0054 | 0.9645 | 0.0057 | 1686.0 | 1961.0 | 1916.5 |

*Florida Museum of Natural History catalog number (http://specifyportal.flmnh.ufl.edu/iz/)

[†]Localities are listed in counter-clockwise geographic order around the state, starting at the panhandle

### GEOGRAPHIC AND TEMPORAL VARIABILITY ASSESSMENT

Assessment of geographic and temporal dimensions of variability in the median ages and corrected posterior age estimates (CPE) for our death assemblage DA samples was complicated by the fact that these metrics vary both stratigraphically and spatially within a given depth interval, meaning the proxy variables available to us (i.e., sample hole and burial depth) are imperfect representations of spatial and temporal variability. Nevertheless, a comparison of sample age and CPE variability with space and depth would be informative about the consistency of the age and time-averaging structure of oyster reef death assemblages.

We used a hierarchical Bayesian model to assess the variability in median age and CPE at the statewide, locality, reef, and sample hole geographic strata. Our model generated locality means from a single statewide mean according to:

$$c_l = e + \zeta_c, l = 1, \ldots, nL, \zeta_c \sim N(0, \sigma_c^2) \qquad (4)$$

where $c_l$ is the locality-level mean at locality $l$, $e$ is the statewide mean, $nL$ is the number of localities, and locality-level means were assumed to be distributed normally with mean of 0 and standard deviation $\sigma_c$. Reef means were generated from each corresponding locality mean according to:

$$r_{kl} = c_l + \zeta_r, k = 1, \ldots, nR_l, l = 1, \ldots, nL, \zeta_r \sim N(0, \sigma_r^2) \qquad (5)$$

where $r_{kl}$ is the reef-level mean for reef $k$ at locality $l$, $nR_l$ is the number of reefs at locality $l$, and reef-level means were assumed to be normally distributed with mean of 0 and standard deviation $\sigma_r$. Observations for the constituent sample holes on each reef were generated from the reef means according to:

$$x_{jkl} = r_{kl} + \zeta_h, j = 1, \dots, nH_{kl}, k = 1, \dots, nR_l, l = 1, \dots, nL, \zeta_h \sim N(0, \sigma_h^2) \quad (6)$$

where $x_{jkl}$ is the observation for sample $j$ from reef $k$ at locality $l$, $nH_{kl}$ is the number of samples from reef $k$, and the sample observations were assumed to be normally distributed with mean of 0 and standard deviation $\sigma_h$. We fit the model to median DA sample age and CPE data for the 15-25 cm and 25-35 cm burial-depths, as well as their difference (25-35 cm values − 15-25 cm values) using the cmdstanr v.0.4.0 interface to Stan in R statistical software v.4.1.1 (Gabry and Cesnovar, 2021; R Core Team, 2021).

The statewide median modeled $e$ for median age was slightly greater for the 25-35 cm burial depth (95 % credible interval of the median difference estimate did not include zero), and there was a much smaller, non-significant, increase in median CPE with burial depth, suggesting that median age tended to increase with burial depth, but the degree of time-averaging did not (Table DR4). In contrast, variation in both median age and CPE tended to increase with burial

TABLE DR4. STATEWIDE RESULTS OF HIERARCHICAL MIXED EFFECTS MODELS OF MEDIAN AGE AND CPE, INCLUDING STATEWIDE MEAN AND STANDARD DEVIATIONS AT LOCALITY, REEF AND SAMPLE HOLE SCALES

| Variable | Category | $e$ | | | $\sigma_c$ | | | $\sigma_r$ | | | $\sigma_h$ | | |
|---|---|---|---|---|---|---|---|---|---|---|---|---|---|
| | | Median | 5 %** | 95 %** | Median | 5 % | 95 % | Median | 5 % | 95 % | Median | 5 % | 95 % |
| Median age | 15-25 cm | 27.90 | 16.66 | 39.55 | 1.26 | 0.16 | 18.03 | 1.99 | 0.25 | 32.47 | 53.14 | 41.81 | 63.36 |
| | 25-35 cm | 40.63 | 18.70 | 64.37 | 0.90 | 0.02 | 7.95 | 74.11 | 3.18 | 100.24 | 28.83 | 22.00 | 79.94 |
| | Difference† | 14.83 | 2.28 | 27.04 | 1.06 | 0.13 | 8.61 | 1.05 | 0.11 | 9.97 | 58.05 | 49.00 | 69.99 |
| CPE* | 15-25 cm | 24.35 | 12.02 | 36.66 | 1.32 | 0.13 | 21.50 | 4.33 | 0.23 | 37.77 | 53.88 | 41.49 | 65.28 |
| | 25-35 cm | 29.06 | 12.00 | 45.15 | 1.08 | 0.12 | 12.96 | 0.93 | 0.11 | 6.24 | 84.47 | 71.54 | 101.32 |
| | Difference† | 6.34 | -11.03 | 23.79 | 1.06 | 0.14 | 14.33 | 0.94 | 0.12 | 7.82 | 96.16 | 81.47 | 114.67 |

*CPE = corrected posterior age estimate

†Differences calculated as (25-35 cm value - 15-25 cm value) for each sample hole

**5 % and 95 % columns denote the 95 % credible intervals

depth (Table DR4, Figures DR2 to DR7). For most variable and category combinations, standard deviations from the locality level and the reef level were similar, but the standard deviations of all combinations increased from the reef level to the sample hole level. Overall, the magnitudes of the variation in differences with burial depth for median age and CPE were comparable to their variation within each burial depth (Table DR4, Figures DR2 to DR7).

These results suggest that spatial variation needs to be considered when planning geochronological investigations of oyster death assemblages because it cannot be assumed that samples taken from the same burial depth within a reef (even only a few meters apart) were deposited contemporaneously. Still, as noted in the main text, comparison of our statewide oyster death assemblage results with those of Dominguez et al. (2016), whose study examined age and time-averaging of death assemblages at six sites in Sydney Harbour (all at ~9 m water depth), suggests that despite the substantial geographic variability in median ages and CPE in our study, the *C. virginica* death assemblages were still more spatially and temporally consistent than a non-reef nearshore shelf molluscan death assemblage (Figures DR2 to DR7).

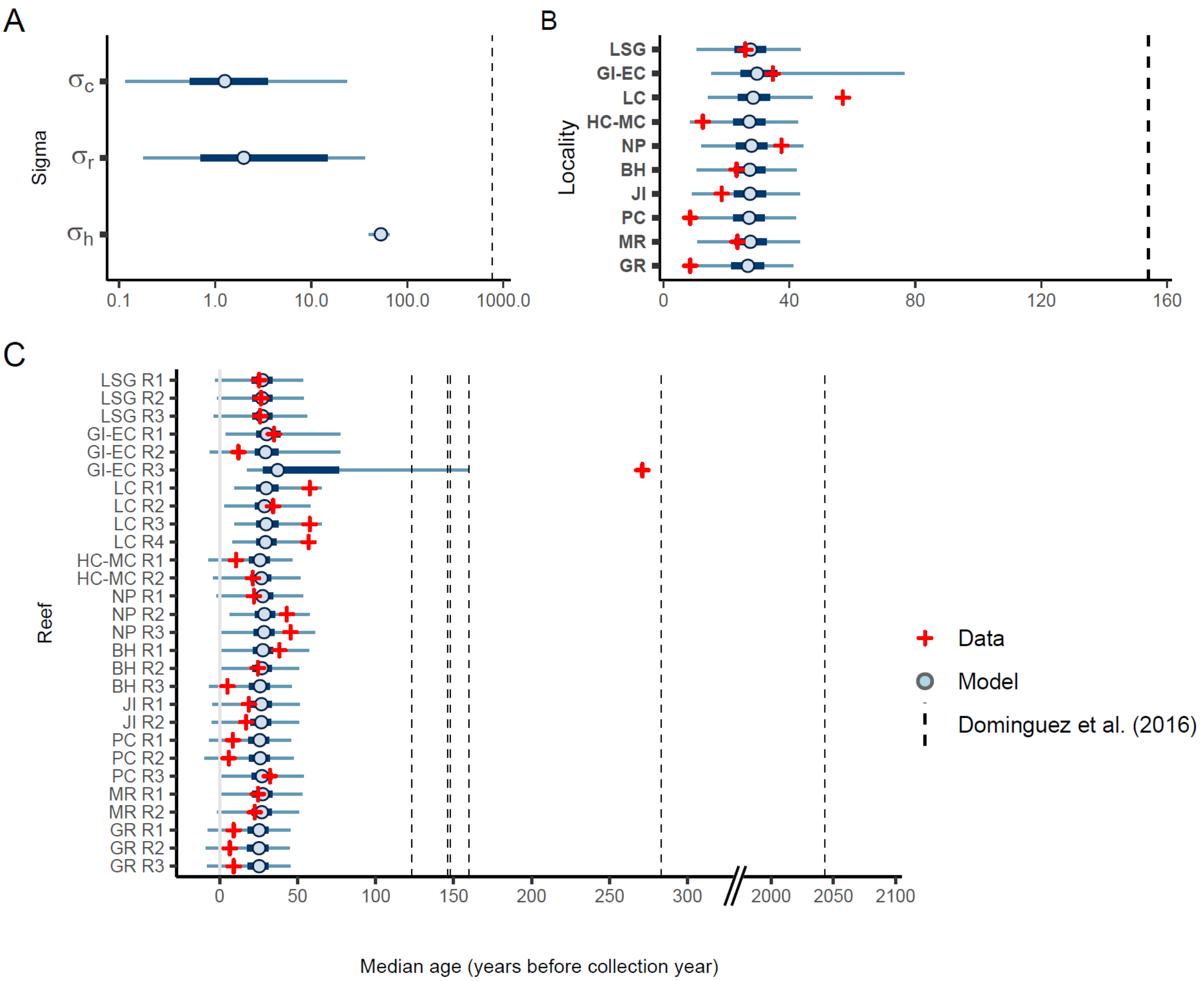


Figure DR2. Plots showing A) estimated standard deviations and medians of the median calibrated ages (relative to 2019) B) by locality and C) by reef for 15-25 cm burial depth in relation to the values calculated from data. Standard deviation, median, and sample-level median ages (relative to 2013) from Dominguez et al. (2016) are also shown in A) to C), respectively, as a comparison between the oyster reef death assemblages and an example of a non-reef (*Fulvia tenuicostata*) death assemblage. Sigma categories correspond to the hierarchical model coefficients (see text for details): $\sigma_c$ = locality-level standard deviation, $\sigma_r$ = reef-level standard deviation, $\sigma_h$ = sample-hole-level standard deviation. Localities are listed on the y axis in counter-clockwise geographic order around the state, starting at the panhandle: LSG = Little St. George Island, GI-EC = Goose Island/East Cove, LC = Lone Cabbage, HC-MC = Hendry Creek/Mullock Creek, NP = New Pass, BH = Big Hickory, JI = Jack Island, PC = Pellicer Creek, MR = Matanzas River, GR = Guana River. Thick horizontal blue lines = 50 % credible intervals, thin horizontal blue lines = 95 % credible intervals.

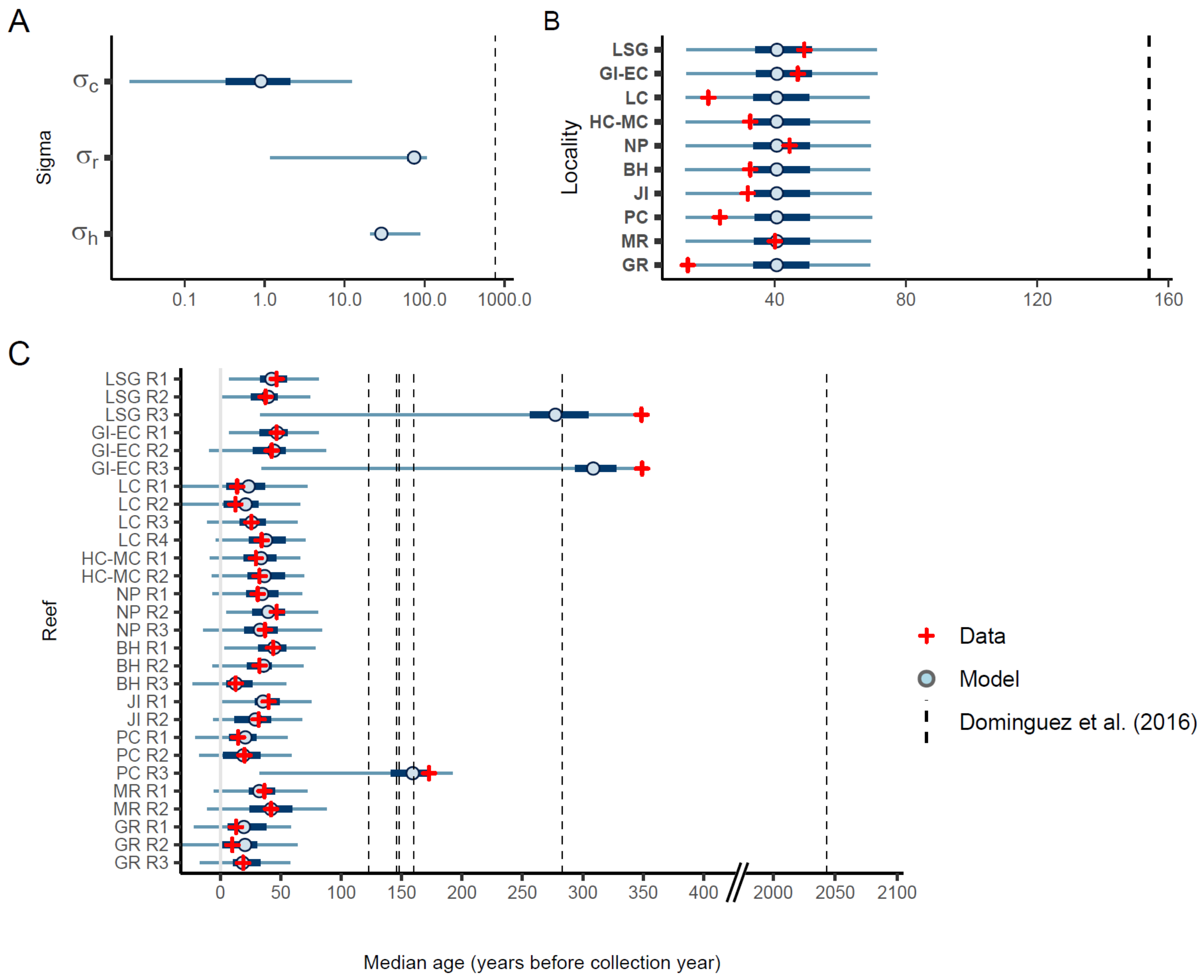


Figure DR3. Same plots as shown in Fig. DR2, but for the DA samples from the 25-35 cm depth interval. See Fig. DR2 caption for plot annotation details.

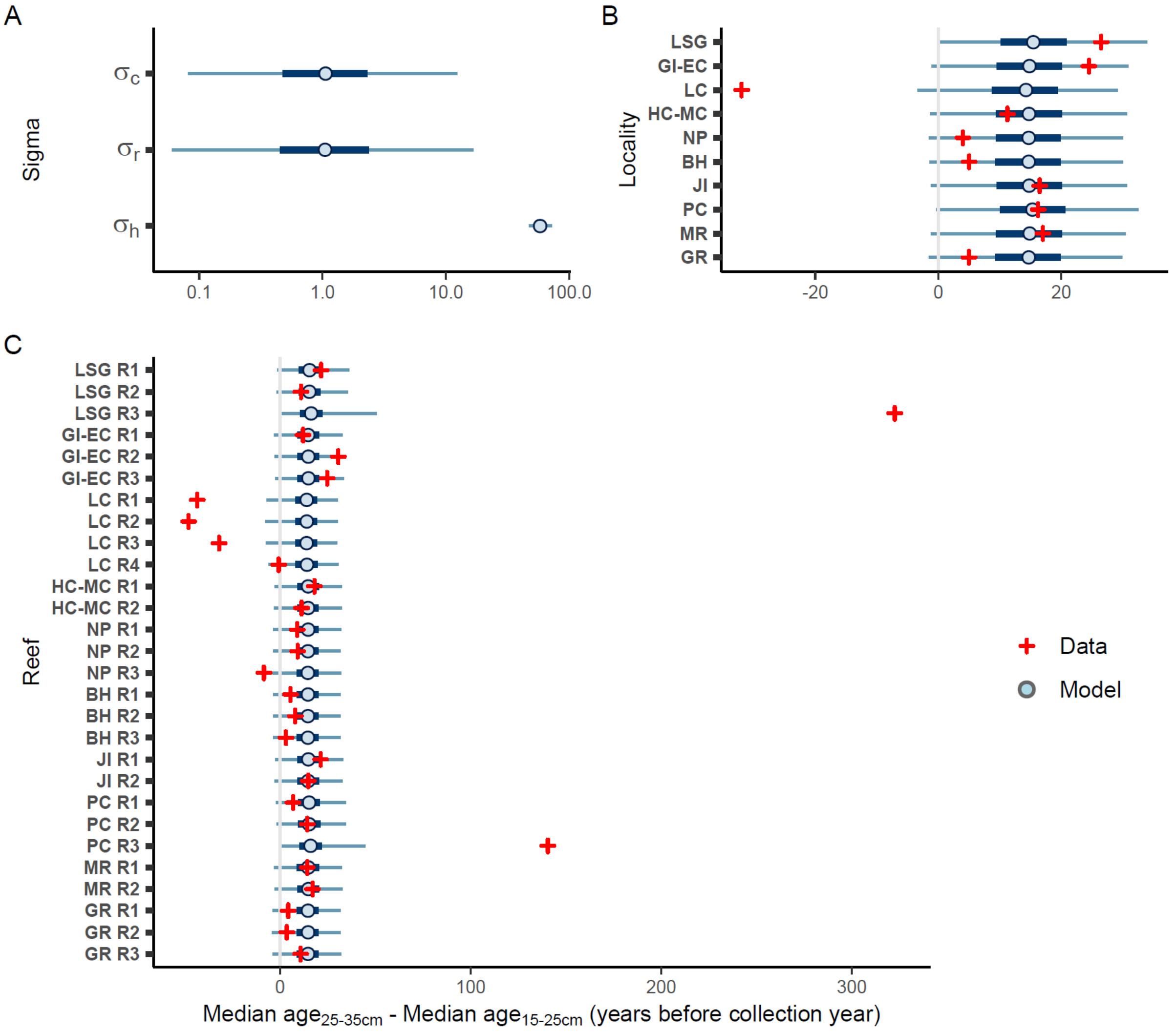


Figure DR4. Plots showing A) modeled standard deviations for locality, reef, and sample hole, and median values for the B) locality and C) reef-level coefficients as in Figs. DR2 and DR3, but for the differences between 25-35 cm and 15-25 cm burial depth median posterior ages. See Fig. DR2 caption for plot annotation details.

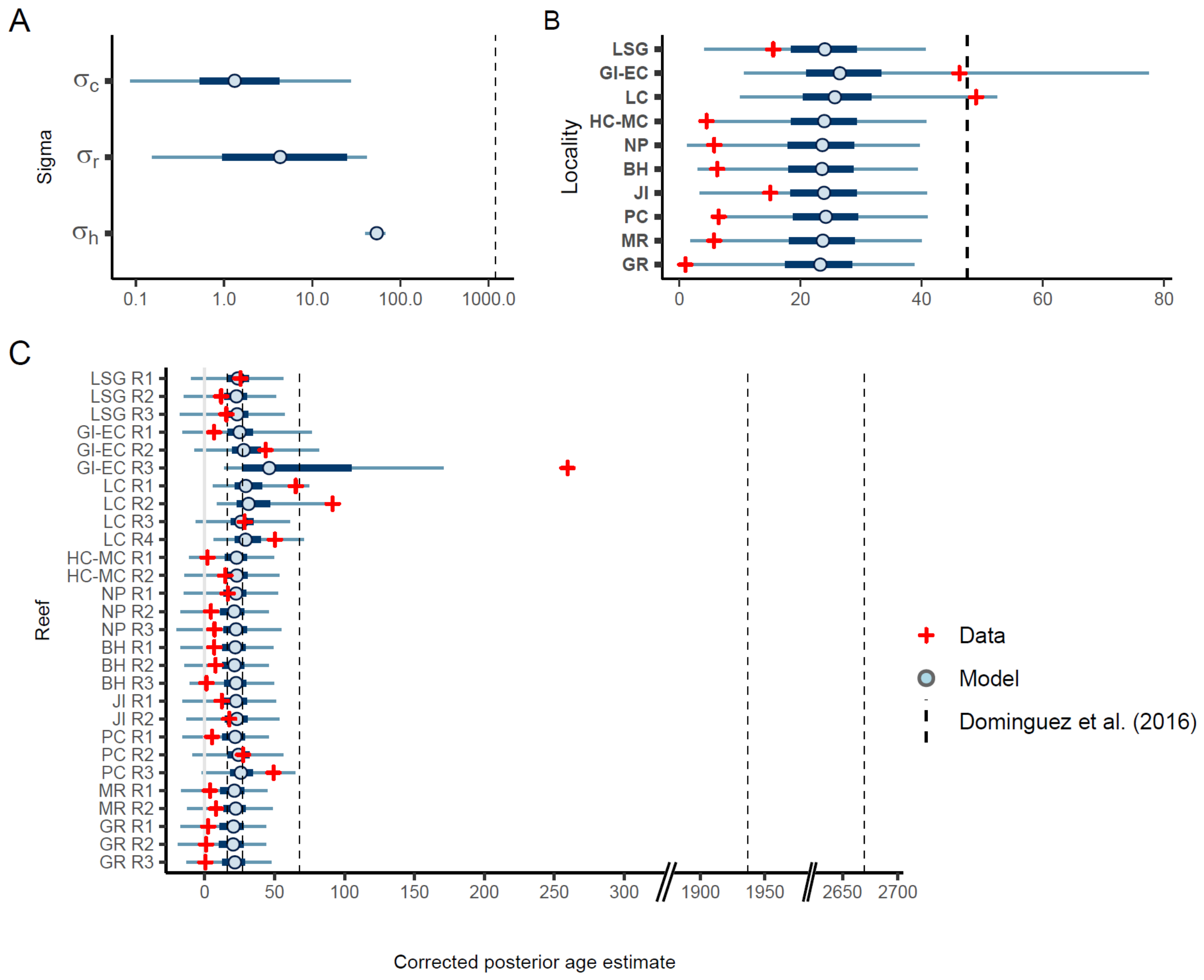


Figure DR5. Plots as described in the caption for Fig. DR2 but showing model results for estimated standard deviations and median corrected posterior age estimates (CPE). See Fig. DR2 caption for plot annotation details.

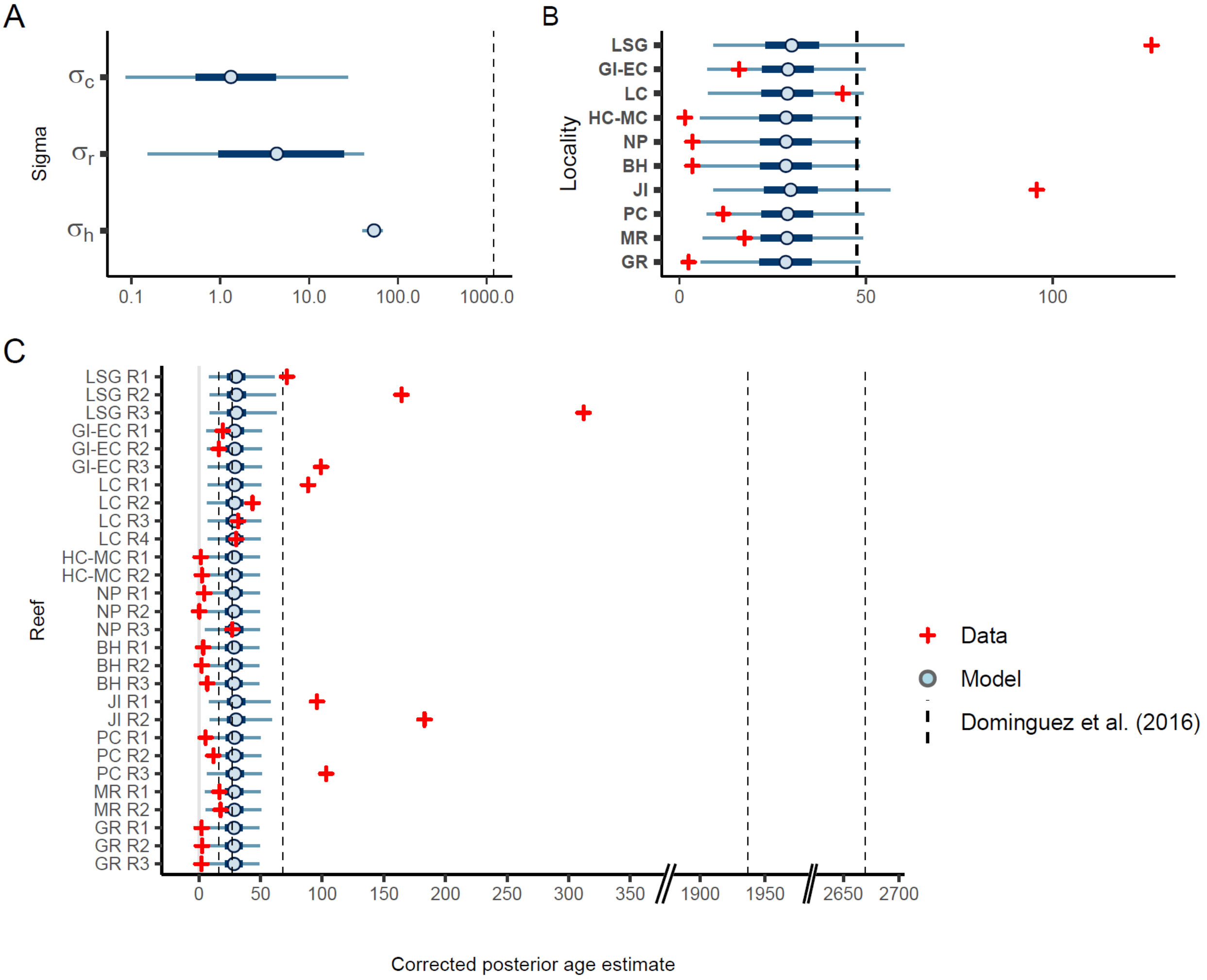


Figure DR6. Plots as described in the caption for Fig. DR3 but showing model results for estimated standard deviations and median corrected posterior age estimates (CPE). See Fig. DR3 caption for plot annotation details.

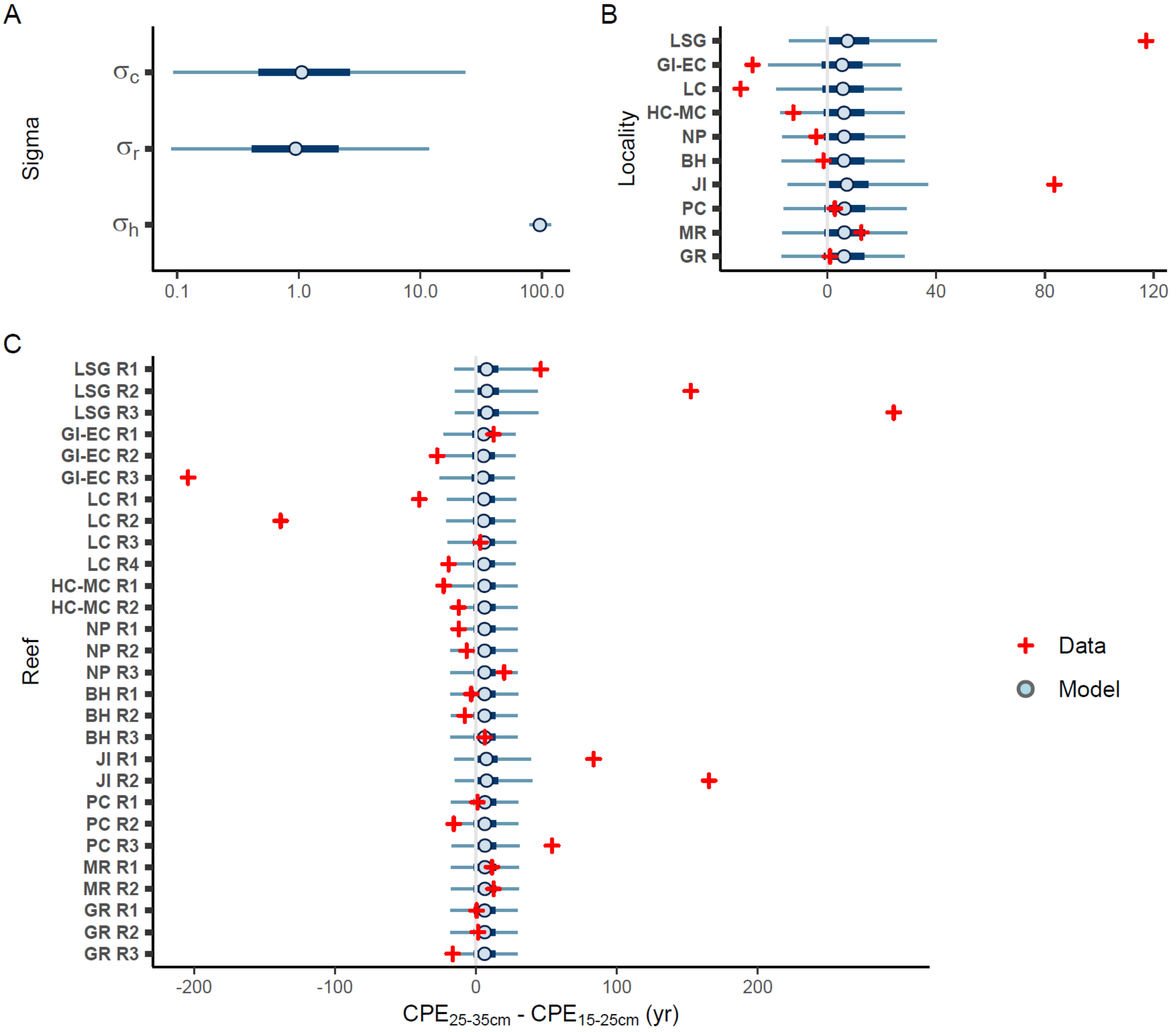


Figure DR7. Plots as described in the caption for Fig. DR4 but showing model results for estimated standard deviations and median differences between 25-35 cm and 15-25 cm burial depth corrected posterior age estimates (CPE). See Fig. DR4 caption for plot annotation details.

**Example of geographic variability in death assemblage characteristics**

Age-depth relationships and scales of time-averaging within an oyster reef DA are products of a complex interaction of processes that can vary on a local scale, including sedimentation rate, reef subsidence, rates of physical and chemical shell destruction on the reef, rates of shell mixing on the reef from storms or bioturbators such as stone crabs, as well as population demographics and recruitment dynamics of the living oyster population, which controls the addition of new shell to the assemblage (Bahr and Lanier, 1981; Hargis and Haven, 1999; Powell et al., 2012; Rodriguez et al., 2014).

These factors are spatially heterogeneous, even on fine spatial scales (i.e., meters; Fig. 2). Some individual reefs in our study showed multi-decadal or even centennial-scale variation in median ages and/or time-averaging estimates for DA samples from the same burial depth. For instance, the minimum and maximum median ages among the three 15-30cm depth interval samples from Reef 1 at New Pass differed by 17 years and the minimum and maximum CPE for the same group of samples differed by 25 years. The overall average within-burial-depth difference between minimum and maximum median DA sample ages by reef (± S.D.) was 24.9 ± 56.8 years, and the corresponding average difference for CPE was 47.5 ± 84.0 years.

Some of the impacts—such as variability in burial rates—are evident in the geochronological results. For instance, DA samples from both burial depths at the Guana River locality are younger than most other localities (Fig. 2). These data are consistent with field observations that suggested a relatively rapid shell burial rate: many Guana River reefs had high relief (~ 1 m), vertically oriented oyster clump growth, and were covered with fine sediments. These characteristics contrasted with those of reefs at other localities, many of which had lower reef heights, coarser, firmer sediments and more rounded, dense oyster clump growth than the Guana River reefs. Altogether, these observations suggest that the burial rate of oyster shell on the reefs at Guana River is more rapid than at reefs elsewhere in the state.
